# Invariant chain with an AP3 interacting sorting signal is sorted to late endosomal compartments and may improve MHC class I loading and presentation

**DOI:** 10.1101/2020.05.13.091579

**Authors:** Ana Kucera, Nadia Mensali, Niladri Busan Pati, Else Marit Inderberg, Marit Renée Myhre, Tone Fredsvik Gregers, Sébastien Wälchli, Oddmund Bakke

**Author notes:** Corresponding authors: Oddmund Bakke, and Sébastien Wälchli.

## Abstract

Invariant chain (Ii) is traditionally known as the dedicated MHCII chaperone. Recent reports have broadened our understanding about various tasks that Ii plays including its physiological role in MHCI cross-presentation. Ii bound MHCI via the MHCII scaffolding CLIP peptide may facilitate MHCI trafficking to the endosomal pathway. The sorting function of Ii depends on two leucine-based sorting signals present in the cytoplasmic tail that acts as binding sites for the adaptor proteins AP-1/AP-2. Here we increased the Ii cross-presentation potency by replacing these with an AP3 motif resulting an efficient transport of Ii from TGN to late endosomes. We also replaced the CLIP region of li with a therapeutically relevant peptide, MART-1. We found the Ii AP3mutant-MART1 construct was capable of loading MHCI and stimulate specific T-cell response more efficiently than the wild type counterpart. The results show that Ii with an AP3 binding sorting motif carrying peptide epitope(s) can promote efficient antigen presentation to cytotoxic T cells (CTLs) independent of the ER located classical MHCI peptide loading machinery.

## Introduction

Vaccination demands successful immune response that in turn depends on activation of both CD8_+_ and CD4_+_ T cells in the context of major histocompatibility complex I and II (MHCI and antigen presentation. The processing of the antigen and its subsequent presentation on MHCI or MHC II at the surface of antigen presenting cells (APCs) requires the coordinated action of different accessory molecules and chaperones. Many strategies have been described to obtain a robust immune response, such as the use of carrier proteins to improve peptide loading to MHC molecules, and targeting of antigens to “favorable” intracellular pathways where MHC reside _1-7_. MHCI and MHCII are not loaded by the same cellular machinery, and they are dependent on different trafficking signals. A successful MHC II antigen presentation largely depends on Ii and peptide loading in the endosomal pathway, while MHCI peptide loading is independent of Ii and occurs primarily in the endoplasmic reticulum _8_.

The targeting to the endosomal pathway of MHCII relies on the two-leucine-based sorting signals Leu_7_/Ile_8_ and Met_16_/Leu_17_ of Ii _9-11_. These signals sequences are present in the cytoplasmic tail of Ii and bind the adaptor proteins AP-1 and AP-2 _12, 13_. These APs are generally found at the trans-Golgi network (TGN) (AP1) and plasma membrane (AP2) and act as coat proteins that bind the donor membrane in order to assemble a scaffold for vesicle budding _14_,_15_. The APs thus mediate sorting from the TGN to endosomes directly or via the cell surface. Upon entry into the endosomal pathway, Ii is sequentially degraded leaving the class II-associated Ii peptide (CLIP) bound to the MHCII groove. CLIP is subsequently exchanged for specific antigenic peptides in the later parts of the endosomal pathway prior to transport to the cell surface for presentation to CD4_+_ T cells. Several vaccination studies have shown that replacement of the CLIP region with an antigenic peptide can lead to efficient MHCII loading and specific presentation to CD4_+_ T cells _1, 2, 16_.

The classical view is that MHCI encounters its (endogenous) antigenic peptides in the ER and this complex is transported to the PM and presents the antigen to the cytotoxic T cell _17_. However, it was demonstrated a few years ago that MHCI like MHCII may be loaded in the endolysosomal pathway guided by Ii _18_, but it was not shown where in the endosomal pathway this took place. An Ii-MHCI interaction was also demonstrated by van Luijn and coworkers who showed that CLIP efficiently binds to several MHCI molecules in leukemic cells _19_. Furthermore, our recent study showed an Ii with CLIP substituted by a MHCI specific tumor antigen is efficient enough to load MHCI and activate T cell specific response in a proteasome/TAP/tapasin independent manner _20_. This strategy was found to be as efficient as exogenous loading of synthetic peptide *in vitro*, and thus identified a novel loading pathway for MHCI which may lead to novel vaccine strategies. In one of our earlier study, we have shown that it is possible to redirect a fusion protein with the li tail to late endosomes by introducing this AP-3 binding motif to the cytoplasmic tail of the li _21_. Here we show that the introduction of AP-3 binding motif in Ii itself re-routed this molecule to proteolytic late endosomal compartments skipping the conventional trafficking route via the PM. As a consequence, this AP3 containing Ii protein had a dramatically shorter half-life than its wt counterpart. We have further investigated the potency of Ii-MHCI mediated antigen presentation with the Ii trafficking mutant designed to bind AP-3 _21, 22_ where we replaced the CLIP region with the cancer relevant peptide MART-1 _20, 23, 24_. This mutant with CLIP/Mart-1 replacement was found to have an improved potency to activate CD8_+_ T-cells compared to the Iiwt and was able to increase the amount of peptide-MHCI on the plasma membrane (PM). Taken together, we find that an improved Ii-based antigen loading/presentation of this peptide may be achieved by routing the Ii and most likely also MHCI to a late stage of the endosomal pathway.

## RESULTS

### Biochemical characterization of IiR_4_RP_6_/L_17_A

The Iiwt sorting signal (Q_4_RD_6_)L_7_I, found to bind the adaptors AP-1 and AP-2 _13_, was replaced as shown in Figure 1A by an (R_4_RP_6_)L_7_I motif, which is a strong AP-3 binder _22_ and found to mediate direct TGN to endosomal sorting _21_. We first tested whether the invariant chain with the AP3 motif, IiR_4_RP_6_/L_17_A was a trimer. In addition to Ii wild type we included the double leucine mutant IiL_7_A/L_17_A known to accumulate at the cell surface due to inactive sorting signals _25_. As shown, all constructs were able to form trimers suggesting that the cytosolic tail mutations did not affect the trimer assembly (Fig. 1B). The abundance of mutant li trimers even in the reduced sample shows the efficiency of the mutant li in making stable trimers that did not compromise on its structural stability (Fig. 1B). However, the protein amount of the AP3 mutant was significantly less than the two others.

**Figure 1:**
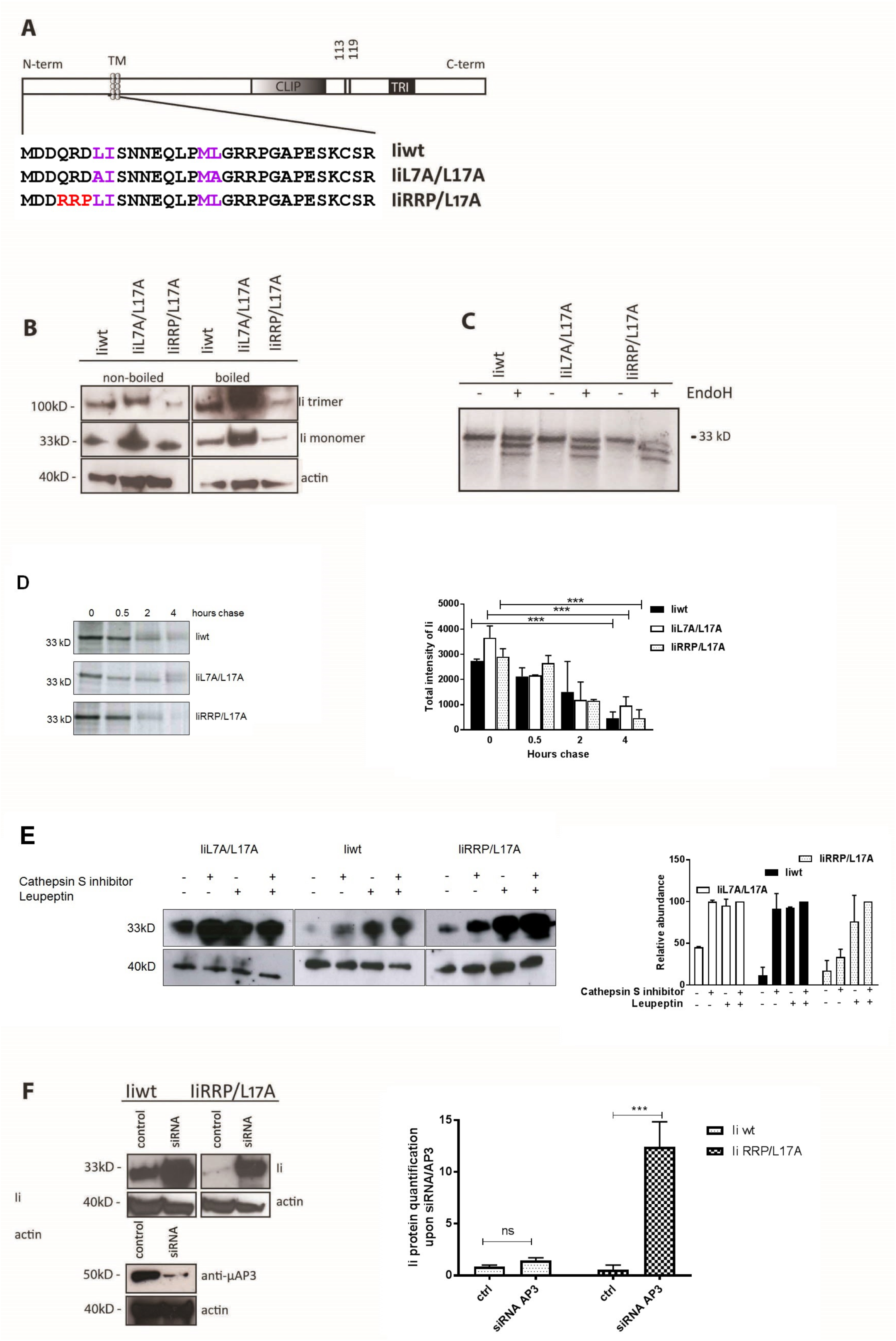
Biochemical characterization of Iiwt and IiR_4_RP_6_/L_17_A. **(A)** Ii constructs used in this study. The mutations done to Ii N-terminal cytoplasmic tail are indicated. CLIP: class II Invariant chain peptide; TM: transmembrane domain; TRI: trimerization domain; position 113 and 119: N-linked glycosylation positions. (**B-E)** For biochemical assays, HEK293 cells were transfected with Ii constructs as indicated. Twenty-four hours later, cleared cell lysates and immune complexes were collected, treated and loaded onto 4-20% SDS-PAGE gels and transferred to PVDF membranes for western blot (WB) analysis. All experiments were repeated three times. (**B)** Lysates were split into two, where half was treated with reducing sample buffer and boiled at 95 °C, while the other was subjected to non-reducing (without β-mercaptoethanol) sample buffer. Lysates were probed with anti-Ii (M-B741) and anti-actin antibodies. (**C-E)** Transfected cells were metabolically labeled with _35_S-Methionine and Cysteine for 30 minutes, washed three times in DMEM and lysed. Radioactivity was detected directly on ECL films. (**C)** Lysates were divided into two where half was subjected to a 15 minutes Endo H treatment prior to immunoprecipitation (IP) with M-B741. Three Ii fractions were found for all constructs; the light /Endo H sensitive ER fraction, the premature Endo H resistant fraction with only one glycan and the fully mature double N-linked glycosylated mature fraction of Ii. **D)** Lysates were collected at time points 0, 0.5, 1 and 4 hours after 30 min of the radioactive pulse, and were immunoprecipitated with M-B741. The radioactivity was further quantified using the tool in ImageJ software, and the graph depicts the protein levels and half-life of the different Ii constructs, as indicated. The data shown in the graph are obtained from three independent experiments. (**E)** After the pulse, cells were incubated for 30 minutes in the presence of: none, one or the other, or both leupeptin and Cathepsin S inhibitor, 24h post-transfection. Lysates were probed with M-B741 and anti-actin. The protein levels were quantified using imageJ. (**F)** HeLa cells treated with either scrambled siRNA (control) or siRNA directed against AP-3, were transfected with Ii constructs as indicated. Lysates were run onto 10% SDS-PAGE gels and transferred to PVDF membranes, where they were probed with anti µ-AP-3, M-B741 and anti-actin antibodies and proteins were quantified using imageJ.

Arginine motifs may mediate ER retention and the Ii mutant (MDDRRPL_7_I) could therefore affect Ii release from the ER thus reducing the total protein level of Ii _26, 27_. To test for this, we performed an Endoglycosidase H (Endo H) treatment, where passage through the Golgi apparatus prior to endosomal sorting is monitored by acquisition of Endo H resistance _28_. Three Ii fractions were detected for the wild type and all the mutants of Ii (Fig. 1C). Thus, all constructs gained Endo H resistance indicating that despite the presence of the RRP amino acid sequence, the Ii mutant can egress the ER. Further to investigate why the level of IiR_4_RP_6_/L_17_A is differed from Iiwt, we performed a pulse-chase experiment to monitor the half-life (t_1/2_) of this protein. Transfected cells were pulsed with _35_Met/_35_Cys containing media, and chased for various time points, followed by an immunoprecipitation of total Ii. As shown in Figure 1D, the t_1/2_ of Iiwt was approximately three hours, whereas IiR_4_RP_6_/L_17_A had a half-life closer to one hour, which suggests a faster kinetic to late endosomal compartments. IiL_7_A/L_17_A shown to accumulate at the plasma membrane _11_ was not degraded after four hours and served as a control. As shown earlier, a TFR fusion protein with the IiR_4_RP_6_/L_17_A cytosolic tail can be targeted directly to the late endosomes/lysosomes _21_. The short half-life indicates that Ii with the IiR_4_RP_6_/L_17_A tail also followed this pathway. To test for such a late endosomal proteolytic localization of IiR_4_RP_6_/L_17_A, we added either the Cathepsin S inhibitor or the broad protease inhibitor Leupeptin, both taken up by endocytosis to the transfected cells. A combination of both cathepsin S and leupeptin were able to protect IiR_4_RP_6_/L_17_A from degradation while the li L_7_A/L_17_A remained protected with either of the protease inhibitors (Fig. 1E). As expected, the Ii wild type, trafficking to the endosomal pathway via the PM _29, 30_ was also protected by both Leupeptin and Cathepsin S. Interestingly, the IiR_4_RP_6_/L_17_A mutant needed both the leupeptin and the cathepsin A inhibitor for maximal inhibition, most likely as this construct is targeted to late endosomes which are more difficult to reach by the endocytosed inhibitors than the wild type Ii which traffics via the PM.

AP-1 is located at the TGN and AP-2 is a plasma membrane adaptor _14_, AP-3 is involved in binding and sorting of proteins to late endosomes and detected both at the TGN and between early and late endosomes and is therefore believed to be involved both in sorting from TGN to late endosomes and endosomal maturation _31_. To further study the pathway of our AP3 binding construct, we performed RNAi depletion of AP-3 and investigated the effect on the protein level. As shown in Figure 1F, AP-3 depletion resulted in a dramatic accumulation of both IiR_4_RP_6_/L_17_A and Iiwt was also protected from degradation, but less. Together with the protective effect of the protease inhibitors in the endosomal pathway, such a strong protective influence of AP-3 depletion on IiR_4_RP_6_/L_17_A is in line with a hypothesis that IiR_4_RP_6_/L_17_A is sorted directly from TGN to the late proteolytic pathway. The control Iiwt is less affected as it is sorted via the PM and only affected by the endosomal maturation inhibition caused by the AP3 inhibition.

### Subcellular distribution of *IiR*_*4*_*RP*_*6*_*/L*_*17*_*A*

The subcellular distribution and trafficking of IiR_4_RP_6_/L_17_A was further investigated using live cell confocal imaging approach. Madine Derby Canine kidney (MDCK) cells were chosen for their high tolerance for laser exposure. The cells were transfected with either Iiwt or IiR_4_RP_6_/L_17_A N-terminally fused with red fluorescent protein mCherry, together with early and late endosomal markers, GFP-Rab5 or GFP-Rab7a respectively. To measure trafficking via the PM the cells were incubated with anti Ii-antibody, M-B741-Alexa647, one hour prior to imaging. It is furthermore known that Iiwt imposes enlargement of endosomes and causes a delay in endosomal maturation _29, 32-34_. Due to this delay, at early time points the antibody reach primarily early endosomes, but gradually throughout the next 2-4 hours, the antibody was also seen in late endosomes _20, 29_. We also observed enlarged endosomes in the cells transfected with Iiwt (Fig 2A and B) and colocalization with GFP-Rab5 and GFP-Rab7a, distributing almost equally (55-60%) within early and late compartments. In addition, more than 60% of Iiwt found to colocalize with M-B741-Alexa 647 (Fig. 2C), confirming trafficking via PM.

**Figure 2:**
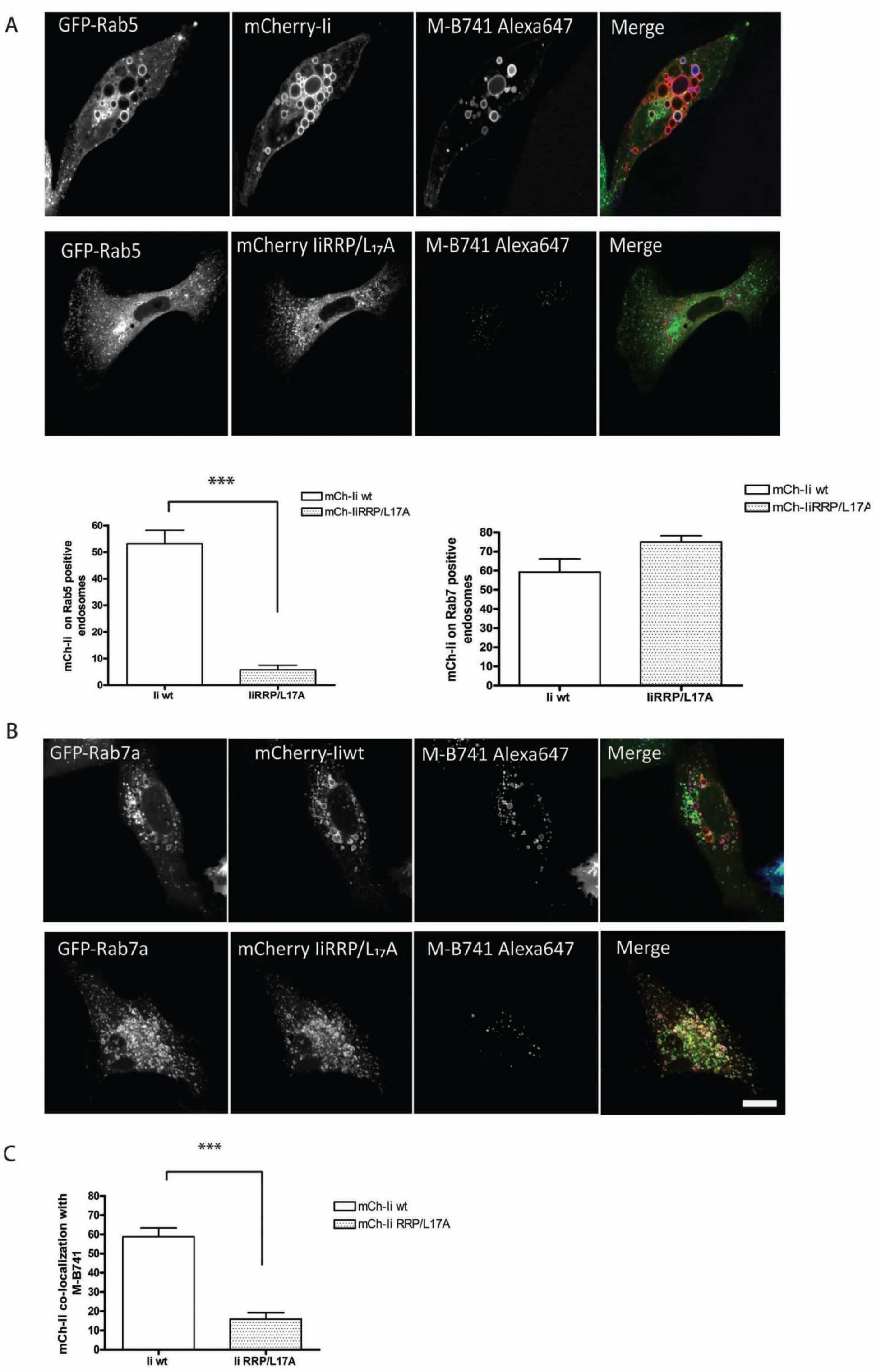
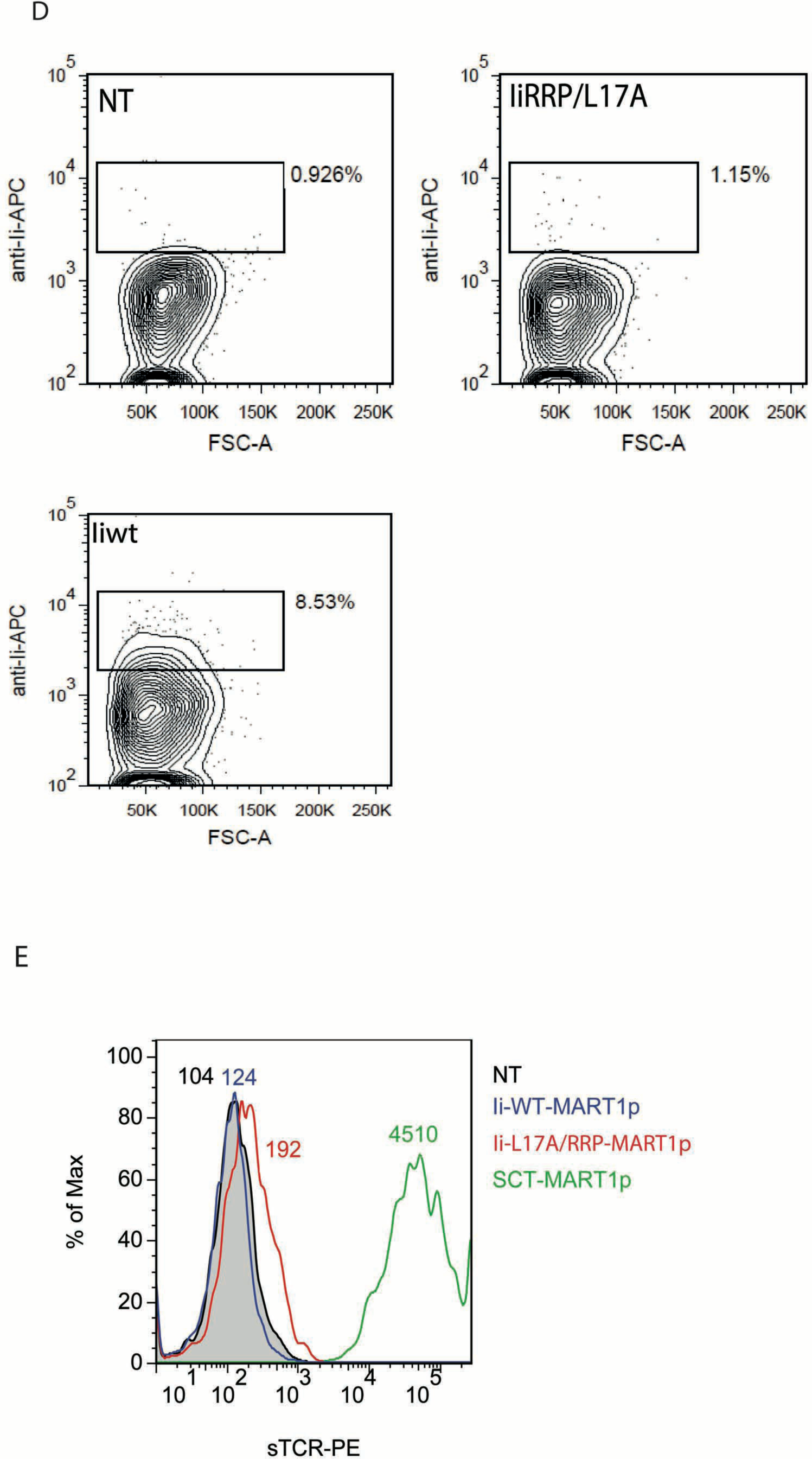
Intracellular distribution of Iiwt and IiR_4_RP_6_/L_17_A. **(A)** MDCK cells were transfected with mCherry-Iiwt or mCherry-IiR_4_RP_6_/L_17_A in combination with GFP-Rab5 or (**B)** GFP-Rab7a as indicated. 24 h post-transfection, the cells were incubated with M-B741-Alexa 647 (anti Ii antibody) for 1h prior to the live cell imaging at 37°C using the Olympus FluoView 1000 inverted microscope equipped with Plan/Apo 60/1.10 NA oil objective. All samples were analyzed in media without phenol red. The images show representative cells and were processed using ImageJ. Three independent experiments were performed, in total 15 cells per condition. Scale bar 15 µm. The graph shows the quantification of the co-localization of Ii in GFP-Rab5 positive vesicles and is shown as mean + SEM of five different cells and indicated *p*-value was determined by unpaired *t*-test *** *P* < 0.001. (**C)** The graph shows the quantification of M-B741-Alexa 647 uptake by the Ii constructs, as indicated, and is shown as mean + SEM of five different cells and indicated *p*-value was determined by unpaired *t*-test *** *P* < 0.001. This graph is representative of three independent experiments, where in total 15 cells were analyzed using ImageJ software. (**D)** SupT1 cells were transduced with Iiwt or IiR_4_RP_6_/L_17_A constructs or left untreated (NT) and stained without permeabilization with anti-Ii antibodies. Presence of membrane Ii was detected by Flow cytometry. This staining is representative of two separate experiments. (**E)** SupT1 (HLA-A2+) cells were transduced with the indicated constructs. Around 10^5^ cells were incubated with FLAG-tagged sTcR (estimated at 10 ng/mL) for 30 minutes at RT. Anti-FLAG antibody was finally used to detect sTcR and the binding was analyzed by flow cytometry. This experiment was repeated once with similar result.

In contrast, mCherry-IiR_4_RP_6_/L_17_A showed a 75% colocalization with the late endosomal marker GFP-Rab7a and less than 10% with GFP-Rab5 (Figure 2A and B). This cellular distribution indicated that the Ii mutant followed a direct sorting route to late endosomes circumventing the cell surface. In further support, we demonstrate that only 12% of mCherry-IiRRP/L_17_A colocalized with M-B741-Alexa 647 (Fig. 2C), which corroborates direct sorting to late endosomes, and is in line with our biochemical characterization of IiR_4_RP_6_/L_17_A. Because of its rapid turnover (Fig. 1D), and the accumulation in late endosomal compartments, IiR_4_R_6_P/L_17_A, did not delay endosomal maturation. The residual uptake of M-B741 was most likely due high expression level of the mutant Ii in some of the cells being missorted to the PM. We confirmed our observations with Ii transduced SupT1 cells. In this assay, the Ii membrane expression was monitored by staining cells without prior permeabilization and later analyzed by flow cytometry (Fig. 2D). This demonstrated that the Ii mutant did not appear on the cell surface. Finally, we used live confocal imaging and confirmed that the IiR_4_R_6_P/L_17_A mutation did not affect the previously described MHCI-Ii association _20_(Sup Fig. 1).

### Soluble T cell receptors detect peptide loaded HLA-A2 from IiR_4_RP_6_/L_17_A

Since the IiR_4_RP_6_/L_17_A mutant is re-routed to a degradation trafficking pathway, we tested whether such a CLIP replaced Ii construct combined with this tail mutation could still load MHCI. To this end, we first detected that the peptide-MHC complex of cell expressing different constructs with a soluble T cell receptor (sTCR) _35_ specific for MART1p. sTCRs have a low affinity for their target, however, we were able to detect Ii-MART1p, inserted in the CLIP region of IiR_4_RP_6_/L_17_A (Fig. 2E). Although very low, this result suggests that the peptide was well loaded on the MHCI molecule. As a control, cells expressing HLA-A2 single-chain trimer (SCT) combined with MART-1 peptide (SCT-M1) were used and showed an expected saturating signal. Taken together, our data support an improved antigen loading ability of IiR_4_RP_6_/L_17_A over the Ii wild type construct. In addition, the trafficking of Ii to the plasma membrane does not seem to be required to get an efficient loading.

### Cells expressing Ii carrying tumor-associated epitopes efficiently load HLA-A2 and specifically activate CD8_+_ T cells

We have previously shown that Ii interacts with MHCI, and that the human MHCI allele, HLA-A*02:01 (HLA-A2) colocalized with Ii throughout the endosomal pathway. We have also demonstrated that CLIP-replaced Ii efficiently activated antigen specific CTLs when expressed in HLA-compatible APC _20_. We therefore compared the ability of the IiR_4_RP_6_/L_17_A mutant to load HLA-A2 peptides with the CLIP-replaced Ii construct (Fig. 3A). J76 cells stably expressing MART-1 specific TCR (DMF5) _35_ were incubated with HLA-A2 positive cells expressing different Ii constructs. IL-2 secretion was used as a read-out for specific TCR stimulation; SCT-MART1 and SCT with an irrelevant peptide (SCT-irr) expressing cells were used as controls. When MART-1 peptide was loaded on HLA-A2 utilizing IiR_4_R_6_P/L_17_A as carrier, the intensity of the stimulation was almost equal to the stimulation observed with SCT-M1, confirming an increase in peptide loading compared to Iiwt (Fig. 3B). In order to support these data, we performed a DC priming study using autologous donor cells. To this end, we assessed the priming ability of DC transfected with Ii mutant (Ii17R_4_RP_6_/L_17_A MART-1) compared to Iiwt MART-1 or the MART-1 peptide. We found that the mutant li (Ii17R_4_RP_6_/L_17_AMART-1) was significantly more efficient and superior to peptide at priming primary CD8_+_ T cells, whereas IiwtMART-1 seemed to be improved but did not reach significance (p=0,052, Fig. 3C), hence at this stage can be considered equal to peptide loading. In addition the Ii17R_4_RP_6_/L_17_A MART-1 was not significantly superior to IiwtMART-1 (p=0.53, not shown). Taken together we can at this stage only conclude that the new construct is functional in DCs, but we might require to test more peptides before we can reach the same conclusions as in the Jurkat system (Fig. 3B). This is in agreement with our previous data where we show that Iiwt construct performed as efficiently as peptide loaded cells _20_. A possible explanation could be that a mutant Ii and wild type bind differently to MHCI. However, by co-immunoprecipitation experiments we found no difference in MHCI binding to the mutated li (Ii17R_4_RP_6_) as compared to binding to wild type Ii (Sup Fig. 2). Together these data support the proposition that IiR_4_R_6_P/L_17_A improved the loading of peptide placed in CLIP region for MHCI mediated antigen presentation.

**Figure 3:**
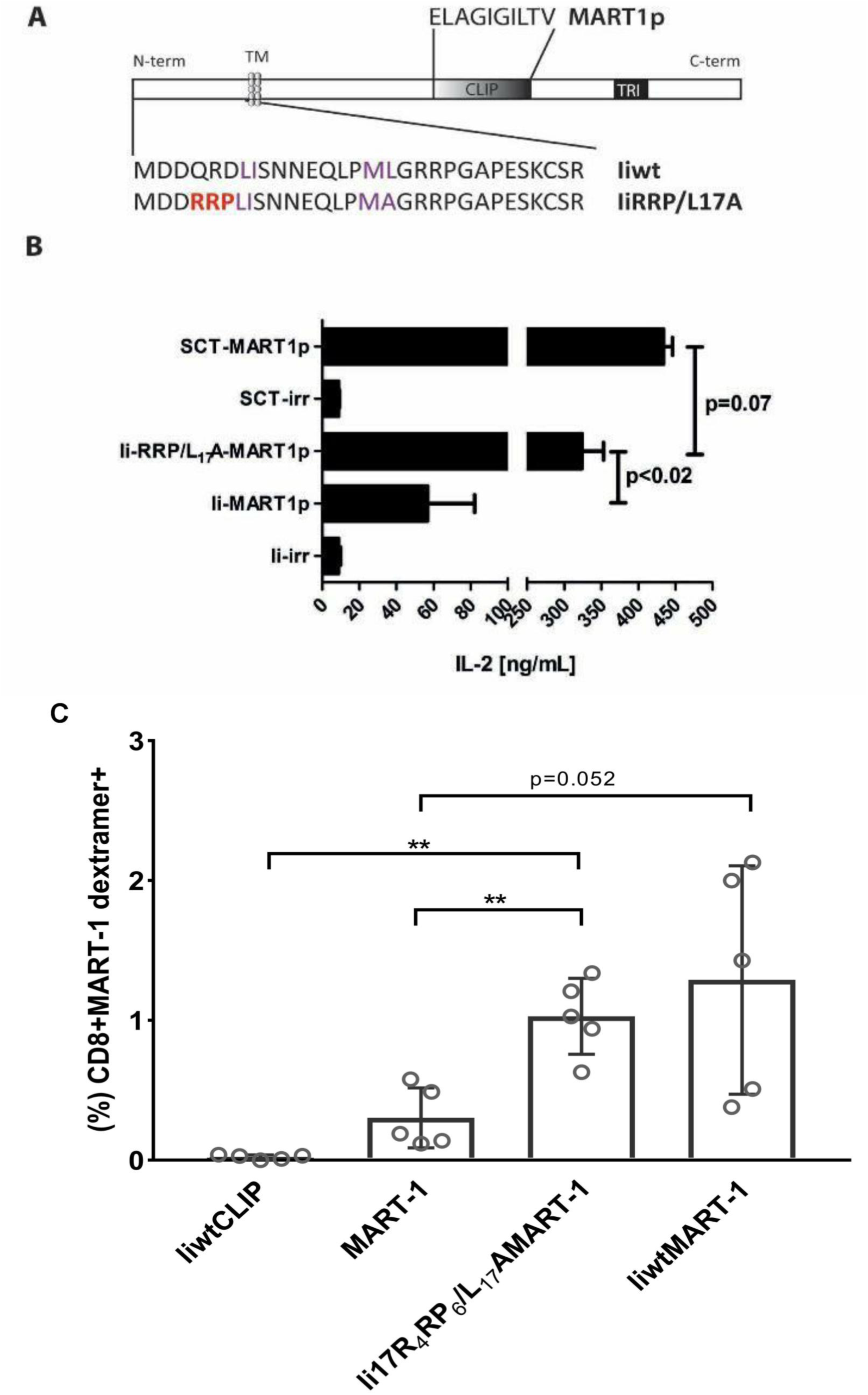
Antigen presentation with trafficking mutant increases MHCI-presentation and naive T-cell priming. **(A)** Ii constructs used in this study. The mutations done to Ii N-terminal cytoplasmic tail are indicated. MART1p: MART1 peptide (ELAGIGILTV); CLIP: class II Ii peptide; TM: transmembrane domain; TRI: trimerization domain. (**B)** SupT1 cells expressing HLA-A2 in combination with the indicated Ii construct (MART1p=MART1 peptide, irr = irrelevant peptide) or a single chain trimer (SCT) construct with MART1p or an irrelevant peptide as controls. J76 cells expressing DMF5 (HLA-A2-MART1p specific TCR) were co-incubated for 20 hours with the presenting cells and IL-2 level in the supernatant was detected by ELISA. The values are given as ng mL_-1_ of IL-2, and error bars are ±SD from two parallels, *p*-value was determined by unpaired *t*-test, this experiment was repeated once with similar results. **(C)** Dendritic cells (DCs) expressing HLA-A2 in combination with the indicated Ii constructs (mutant Ii17R_4_RP_6_/L_17_A carrying MART-1 peptide or wild type li, (liwt), carrying MART-1 peptide) were generated as wells as a wild type li construct with unmodified CLIP region (liwtCLIP) and DCs loaded with MART-1 peptide as controls. Autologous T cells were primed with the presenting cells (DCs) and co-incubated for 8 days. On day 9, T cells were restimulated with DCs and the presence of MART-1 specific T cells was detected by MART-1 dextramer staining 19 days later. The values are given as frequency (%) of CD8+MART-1 dextramer+ T cells and error bars are standard deviations (n=5), *P*-value was determined by unpaired *t*-test with Welch’s correction and analysis was performed by GraphPadPrism software (GraphPad Software, San Diego, CA, US).

## DISCUSSION

Previous studies have revealed that neither MHCI nor MHCII are AP3 dependent for trafficking and antigen presentation _36_. The kinetics of Ii transport and degradation are also unaffected in cells lacking AP-3 _37_. Through introduction of the AP-3, instead of the AP1/AP2 binding motif, we successfully re-routed the Ii towards late endosomal compartments. In addition to the re-routing of Ii, the insertion of AP3 binding motif increased the kinetics of trafficking of Ii to the late endosomal compartment. Overall, our effort of bringing mutations in the Ii created a strict and direct sorting pathway with improved MHCI peptide loading efficacy. Here, we characterized the sub-cellular distribution of IiR_4_RP_6_/L_17_A and compared with the Iiwt. Equal distribution of Iiwt to the early and late endosomes was observed whereas IiR_4_RP_6_/L_17_A was found to be mainly colocalizing with late endosomal markers. The direct sorting to endosomes also overcomes the property of delayed endosomal maturation _29_ substantiating a faster antigen processing.

Recent studies have shown the role of Ii in trafficking of MHCI to the endosomal pathway and its implication in cross presentation _18, 20_. The description of Ii as a vehicle to perform antigen presentation has brought this molecule to the doorstep of the clinic not only as a target for immunotherapy _38_ but also as a vector for increasing immune reactions towards specific oncogenic antigens _20_. Genetic exchange of the CLIP region with a peptide antigen substantially loads MHCI and presents the antigen on the cell surface. Our results showed that the antigenic peptide carried by IiR_4_RP_6_/L_17_A can be detected by T cell carrying a specific TCR. Additionally, the antigen detection by the sTCR supports the specificity and the robustness of the IiR_4_RP_6_/L_17_A mutant loading. Taken together, our data show that the modification in trafficking induced by the IiR_4_RP_6_/L_17_A mutation can improve peptide loading and thus MHCI-peptide levels at the cell surface. In this study, IiR_4_RP_6_/L_17_A carrying MART-1 antigen was shown to be equally efficient in activating specific TCR carrying cells in comparison to liwt. In addition, we found that IiR_4_RP_6_/L_17_A mutant was also competent to load enough peptide onto DC in order to prime primary naive T cells. These studies confirm the capacity of MHCI loading in the endosomal pathway and establish that the loading may take place in the proteolytic later parts of this pathway. Additional studies will be necessary to determine if such and AP3 binding mutation will be advantageous *in vivo*, for instance to improve immunotherapy using modified Ii as an immunization vector.

## METHODS

### Recombinant cDNA constructs

cDNA encoding human Iip33 wt _9_, was subcloned into the pcDNA3 expression vector at *Kpn*I-*Bam*HI. Human Iip33 mutants, Ii L_17_A and Ii L_7_A L_17_A, in the PSV51L expression vector have also been described _25_. *Kpn*I and *Bam*HI restriction sites were introduced up and downstream of the Ii sequences respectively by PCR and the following primers were used: Ii-*Kpn*I forward 5’ AGAGA GGGTACCGTCATGGATGACCAGCGCGAC 3’. Ii-*Bam*HI reverse - 5’ AGAGAGGGATCCTCACATGGGGACTGGGCCCAG 3’. The Ii mutants were thereafter subcloned into pcDNA3 at *Kpn*I-*Bam*HI, behind the T7-RNA polymerase promoter, and Ii L_17_A was subsequently used as template for PCR quick change mutagenesis (all reagents used were included in the kit; QuickChange® Site-Directed Mutagenesis (Stratagene, La Jolla, CA, USA)) in order to generate the AP-3 binding motif RRP _21_. Primers used: L17A RRP sense 5’ CCGTCATGGATGAC<u>CGTCGTCCC</u>CTTATCTCCAACAATG 3’ and L17A RRP anti-sense 5’ CATTGTTGGAGATAAG<u>GGGACGACG</u>GTCATCCATGACGG 3’. All primers were purchased by Eurofins MWG Operon (Ebersberg, Germany). mCherry-Ii was made by cloning Iiwt in frame to the C terminal end of mCherry without the stop codon in pcDNA3 (a kind gift from Terje Espevik, NTNU, Trondheim, Norway). mCherry-IiR_4_R_6_P/L_17_A was purchased from GenScript (Piscataway, NJ, USA). HLA-A2-GFP has been described and transfections were carried out as already described _20_. GFP-Rab5 and GFP-Rab7 were supplied by Cecilia Bucci _39_. All CLIP-antigenic peptide constructs (IiMART1) were cloned by site direct mutagenesis of the Iiwt and IiR_4_RP_6_/L_17_A construct subcloned in pENTR vector (Invitrogen, Oslo, Norway). The mutagenesis to change the CLIP peptide (MRMATPLLM) into antigenic peptides (MART1: ELAGIGILTV) was performed as described _20_. After sequence verification, these constructs were recombined into a Gateway-converted pCI-pA102 _40_

### Cell culture, transfections and RNA interference

HEK293 cells, human epithelial HeLa-Oslo and Madin Darby Canine Kidney (MDCK) cells were grown in Dulbecco’s Modified Eagle Medium (DMEM, Bio Witthaker, Walkersville, MD, USA). All media were supplemented with heat-inactivated 10% fetal calf serum (FCS, HyClone, Logan, UT, USA). J76 were a kind gift from Miriam Hemskerk, (Leiden University Medical Center, Leiden, The Nederland) SupT1 from Martin Pule (UCL, London, Great Britain), both cell lines were grown in RPMI+10% fetal calf serum. All cells were grown in a 5% CO_2_ incubator at 37°C. Transient transfections were performed with either lipofectamine 2000 reagent from Invitrogen (Hek293 cells, MDCK) or with FuGENE 6 (ProMega) (HeLa), both according to manufacturer’s protocols. For siRNA interference (RNAi) we used the following oligonucleotides; the sense µ3A, 5′-GGAGAACAGUUCUUGCGGC-3′ and the antisense 5′-GCCGCAAGAACUGUUCUCC-3′ oligos, for negative control a scrambled sequence was used, sense: 5′ ACUUCGAGCGUGCAUGGCUTT 3′ and antisense scrambled control 5′ AGCCAUGCACGCUCGAAGUTT 3′. All of the oligos were from Eurofins MWG Operon (Ebersberg, Germany) and are previously described _36, 41_. Transfection of HeLa with siRNA was performed as previously described _42_.

### Antibodies and reagents

M-B741 was purchased from BD Biosciences (Franklin Lakes, NJ, USA). Labeling of antibody was with Alexa-647 performed according to manufacturer’s protocol (Invitrogen/Molecular Probes, Carlsbad, CA, USA). Anti-actin was purchased from AbCam, (Cambridge, UK). The anti AP-3 antibody is affinity-purified rabbit antiserum directed at its µ subunit and was a kind gift from Professor Margaret S. Robinson (Cambridge, UK). The secondary antibodies: sheep anti-mouse- and sheep-anti rabbit-HRP were acquired from Invitrogen/Bio-Rad (Hercules, CA, USA). Anti-FLAGM2 monoclonal antibody was purchased at Sigma-Aldrich (Oslo, Norway). Soluble DMF5 TcR was prepared as described by Walseng *et al*. _35_.

### Biochemical analyses

Metabolic labeling was performed using S_35_-labeled Cysteine/Methionine (Perkin Elmer, Waltham, MA, USA). Cells were seeded to 60%-70% confluence in 6-well plates; washed three times in Cys/Met-free DMEM; incubated in Cys/Met-free DMEM for 45 min followed by a 30 minutes pulse with Cys/Met-free DMEM supplemented with 50µCi S_35_. For the pulse chase assay, the cells were washed three times in DMEM containing 2mM L-glutamine, primocin, and 30% FCS and chased for indicated time periods. Immunoprecipitations were done at 4 °C over night with 1-2µg ml_-1_ antibody in lysis buffer (50 mM Tris-HCl, pH 7,5, 150 mM NaCl, 1% Tx100) supplemented with the protease inhibitor cocktail Protease Arrest (G-Biosciences, St. Louis, MO, USA). Antigen–antibody complexes were captured with Protein G-coupled Dynabeads (Invitrogen) and re-suspended in gel loading buffer containing 2% SDS, 125mM TrisHCL, 20% glycerol and 5% β-mercaptoethanol, or its non-reducing β-mercaptoethanol free counterpart. The samples were boiled for 5 min at 95° C, loaded onto 4-20% Tris-HEPES-SDS gels (Pierce, Rockford, IL, USA), and transferred onto PVDF membranes (Millipore, Billerica, MA, USA). Antibody incubation was done in 5% skim milk (BioRad, Hercules, CA, USA) at room temperature and immunoprecipitated protein was detected using Amersham® ECL Plus Western Blot Detection System (GE Healthcare, Buckinghamshire, UK). Radioactivity, however, was detected directly on ECL films (GE Healthcare, Buckinghamshire, UK). For the experiments including protease inhibitors, the same procedure was followed as for metabolic labeling. During the 30 min S_35_-Cys/Met pulse, 20 nM Cathepsin S Inhibitor (Merck Chemicals Ltd., Nottingham, UK) and/or 100 μM Leupeptin (SIGMA ALDRICH) were added. The procedure was then continued as described above. For Endo H digestion, the beads were resuspended in 0.1 M sodium phosphate buffer (pH5.5) containing protease inhibitor as described above. The samples were divided into two and incubated for 15 minutes at room temperature with, or without 0,5 mU of Endo H (SIGMA). After the Endo H treatment, the samples were boiled at 95°C, and loaded onto gels as described above.

### Flow cytometry

Indicated samples were acquired using a BD LSR II flow cytometer and the data were analysed using FlowJo software (Treestar Inc., Tilburg, The Netherlands).

### Spinning disk – and confocal laser scanning microscopy

MDCK cells were grown to 70% confluence in 35 mm microwell dishes (MarTek, Ashland, MA, USA). The cells were then transfected with; GFP-Rab5, GFP-Rab7a, Ii constructs and HLA-A2-GFP/β2m. 1h prior to imaging, the cell medium was exchanged with complete DMEM without phenol red, and cells were incubated with M-B741-Alexa 647 to a final concentration of 1 µg/mL. Live imaging was performed to eliminate fixation artifacts. Confocal images were acquired on an Olympus FluoView 1000 inverted microscope equipped with Plan/Apo 60/1.10 NA oil objective (Olympus, Hamburg, Germany). Constant temperature was set to 37 °C and CO_2_ to 6% by an incubator enclosing the microscope stage. Fluorochromes were exited with 488nm, 543nm and 647nm lasers. All image acquisition was done by sequential line scanning to eliminate bleed-through. Live films were acquired using an Andor Revolution XD Spinning Disc microscope with PlanApo 60×1.42 NA oil immersion objective, as this microscope provides an ideal platform for high speed, high signal to noise imaging, with low bleach rate and low photo-toxicity. Three lasers were used; 488nm, 561nm, and 640 nm, and 4 frames per minute were acquired for the total of 25 min. Images was processed with ImageJ (NIH, USA) and Illustrator (Adobe systems Inc., San Jose, CA, USA).

### *In vitro* generation of Dendritic cells for antigen presentation and T-cell priming assay

Immature dendritic cells (DCs) were generated essentially as described in Subklewe, *et al*. _43_ Briefly, monocytes obtained from leukapheresis product (REC Project no: 2013/624-15) were cultured for 2 days in CellGro DC medium (CellGenix, Freiburg, Germany) supplemented with GM-CSF and Interleukin-4 (IL-4) in Ultra-low attachment cell culture flasks (Corning). The immature DCs were electroporated with either mRNA encoding for mutant Ii17R_4_RP_6_/L_17_A MART-1 or wild type Ii (liwt) carrying MART-1 peptide. Cytokines facilitating maturation were used (IL-1β, IL-6, TNF-α, IFN-γ (all from PeproTech, Rocky Hill, NJ), prostaglandin E_2_ (PGE2), and TLR7/8 agonist R848 (MedChem Express, Sweden)) _44_ and cultured for 24h. Mature DCs were used in T cell priming experiment. DCs electroporated with wild type li mRNA (no CLIP replacement) and DCs loaded with MART-1 peptide (10 µM) were included as negative and positive control, respectively, in the priming assay. Briefly, the distinct DC populations were cultured with autologous PBMCs at 1:10 DC:T cell ratio. On day 3, T cell cultures were supplemented with IL2 and IL7.On day 8, T cell cultures were re-stimulated with DCs and 10 days later T cells were stained with MART-1 dextramer (Immudex, Copenhagen, Denmark) to assess the presence of MART-1 antigen-specific T cells in the cultures.

## ACKNOWLEDGEMENTS

We thank the NorMIC Oslo imaging platform at the Department of Biosciences, University of Oslo for use of imaging facilities. The financial support of the Norwegian Cancer Society (grants 4604944 to O. B.), the Research Council of Norway (grant 230779 to O. B., grants 244388 and 254817 to E.M.I and M.R.M. and N.M, respectively and through its Centre of Excellence funding scheme to O.B., project number 179573) is gratefully acknowledged and South-Eastern Norway Regional Health Authority to S.W. (Innovation grant 13/00367-88) and to E.M. (grant 2010021).

