## Supplemental figures for "Invariant chain with an AP3 interacting sorting signal is sorted to late endosomal compartments and may improve MHC class I loading and presentation"

### Figure Sup. 1

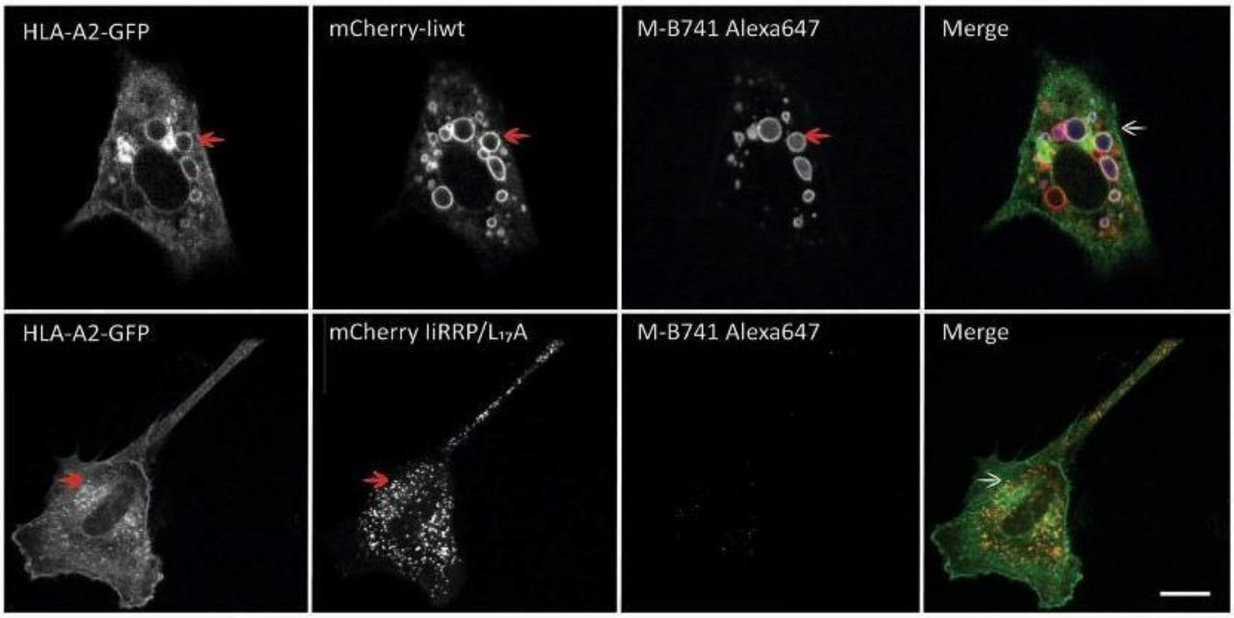


**IiRRP/$L_{17}$A colocalizes with HLA-A2-GFP in the endosomal pathway**

MDCK cells were transfected with HLA-A2-GFP, mCherry-Iiwt and mCherry-$R_4RP_6/L_{17}A$ as indicated, and 24h post-transfection, the cells were added M-B741-Alexa 647 for 1h prior to live cell imaging at 37°C using the spinning disk confocal microscope with a 60x/1.42 NA oil immersion objective. All samples were analyzed in microscope media without phenol red. Images were acquired every 15 sec for 20 minutes. The experiment shown was repeated thrice, and representative films are shown. Scale bar 10µm.

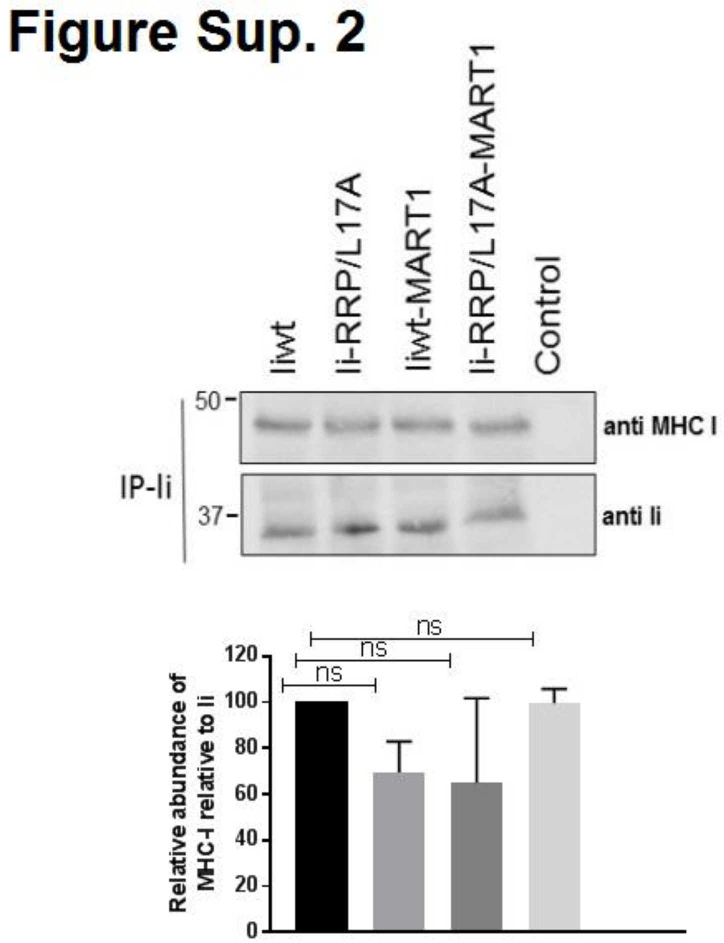


**MHC-I does not preferentially bind to mutant Ii**

MDCK cells were transfected with the indicated Ii constructs along with human MHC-I. After 24 h cells were lysed and Ii was immunoprecipitated with M-B741 bound to protein G-dynabeads. Co-immunoprecipitated MHC-I protein was detected by using anti-human MHC I antibodies. The same membrane was also stained with anti-human Ii antibodies. Non-transfected MDCK cells were used as a control. The band intensity of MHC-I in relative to Ii is quantified using imageJ software. The data shown here are the result of two independent experiments. ns: not significant.
